# Polygenic and monogenic adaptation drive evolutionary rescue at different magnitudes of environmental change

**DOI:** 10.1101/2025.06.13.659553

**Authors:** Tatiana Bellagio, Moises Exposito-Alonso

## Abstract

Understanding the genetic basis of rapid adaptation is key to predicting species’ evolutionary responses to environmental change. However, it is still debatable whether many small-effect mutations or a few large-effect mutations underlie rapid adaptation, and how this knowledge can predict population survival or extinction. To address this question, we performed a series of ecologically grounded forward-in-time genetic simulations to study rapid adaptation and extinction with increasing magnitudes of environmental change. These simulations were seeded with genomic variation of the plant *Arabidopsis thaliana* to have a realistic genomic structure, with one (monogenic) to 1,000 (polygenic) variants with varying heritabilities contributing to an environmental adaptive trait. Our results revealed two distinct scenarios of rapid adaptation and population rescue. Under small to moderate environmental shifts, high polygenic traits increased evolutionary rescue probability. Under extreme environmental shifts, high polygenic traits lead predictably to extinction, yet monogenic traits sometimes produce one-off winning adaptive genotypes. We interpret our rapid evolutionary rescue findings in terms of the fundamental theorem of natural selection, where monogenic and polygenic traits differ in how they create stable versus skewed fitness variance (*V*_*w*_) and how they respond to environmental shifts. These results highlight the insights genomics gives us into the (un)predictability of species’ evolutionary responses to global change, with management implications for assisted adaptation conservation.

## Introduction

When environmental changes displace species from their ecological niches, they must either track suitable conditions through range shifts, adapt to novel environments, or face local to global extinction (Wiens 2016; Parmesan 2006; de Lafontaine et al. 2018). Populations’ adaptive capacity—driven by natural selection acting on within-species genetic variation—has become a critical survival determinant in a rapidly changing world (J. N. Thompson 1998; Barrett and Schluter 2008; IPCC 2022). Contrary to the classical view of evolution as a slow process, empirical studies in the genomics era have now demonstrated that rapid, genetically-based phenotypic changes are widespread. Examples include industrial melanism in peppered moths spreading in less than 50 years (Cook and Saccheri 2013); insecticide resistance in *Drosophila sp*. evolving within a few decades of insecticide application (Daborn et al. 2002); climate-driven shifts in flowering time in *Brassica rapa*, observed over just a few generations (Franks, Sim, and Weis 2007); and widespread herbicide resistance in agricultural weeds, emerging within decades of herbicide deployment (Heap 2024). Critically, such evolutionary dynamics may have consequences at the demographic level and, in cases of abrupt or extreme environmental change, may rescue a population from decline and extinction—achieving evolutionary rescue. (Gomulkiewicz and Holt 1995; Bell and Gonzalez 2009; Carlson, Cunningham, and Westley 2014). Assessing the likelihood of within-population genetic diversity catalyzing evolutionary rescue in natural populations plays a key role in applied evolutionary biology for conservation strategies.

Whether populations can adapt depends not only on the presence of within-species genetic variation, but on its genetic architecture: the number, effect sizes, and interactions among adaptive loci across the genome, including complex phenomena such as dominance, pleiotropy, and epistasis. Among these dimensions, a long-standing question in evolutionary biology concerns whether populations typically evolve through the cumulative effects of many small-effect loci or a few large-effect loci (Orr 2005; Connallon and Hodgins 2021). This debate, rooted in early 20th-century genetics, contrasts the Mendelian view that traits are governed by a few large-effect mutations with the biometrician perspective—later formalized by Fisher—that quantitative traits arise from the cumulative effect of many small-effect loci (Fisher 1918; Pritchard and Di Rienzo 2010; Stern and Nielsen 2019). As a theoretical synthesis, Orr proposed that adaptive substitutions follow an exponential distribution of effect sizes, with adaptation driven by both many small-effect and a few large-effect alleles (Orr 1998). Empirical evolutionary studies have identified prominent examples of well-described large-effect loci involved in adaptation, such as *CORTEX* driving industrial melanism in the peppered moth (Van’t Hof et al. 2016; Sheehan et al. 2016) and *YUP* influencing flower color transitions in *Mimulus sp*. (Liang et al. 2023), among many others (Dorweiler et al. 1993; Cooley et al. 2011; S. D. Smith and Rausher 2011). Concurrently, quantitative genomics literature has reported thousands of small-effect loci underlying many important traits across species (Khaipho-Burch et al. 2023). Examples of such traits are human height, where the number of identified loci continues to grow with increasing sample size (Visscher et al. 2017; Boyle, Li, and Pritchard 2017), or grain yield traits in plants (Zaw et al. 2019; Cao et al. 2020). Regardless of the specific genetic architectures of adaptive traits across species, we still lack a clear understanding of when different architectures may benefit or prevent evolutionary rescue in natural populations.

The study of different genetic architectures and their influence on adaptation has typically involved theory and simulations with a focus on adaptation temporal dynamics. Much of the early work, rooted in classical population genetics, focused on understanding how beneficial alleles of large effect sweep to fixation (J. M. Smith and Haigh 1974; Kaplan, Hudson, and Langley 1989; Hermisson and Pennings 2005; Barrett and Schluter 2008). In contrast, work on polygenic adaptation was initially studied through the infinitesimal model, which treats traits as continuous and does not explicitly track individual loci (Lande 1976; Walsh and Lynch 2018). More recent theoretical and simulation-based studies have modeled shifts in allele frequencies across many small-effect loci, increasingly incorporating biologically realistic factors such as demography, effect size distributions, and environmental change (Barton and Etheridge 2018; John and Stephan 2020; Barghi, Hermisson, and Schlötterer 2020; Hayward and Sella 2021; Jain and Stephan 2017). To bridge the gap between monogenic and polygenic adaptation, recent work has explored conditions under which one architecture is favored over the other. These include redundancy and initial allele frequency (Höllinger, Pennings, and Hermisson 2019), the magnitude of environmental change (Milligan, Hayward, and Sella 2025), and selection spatial structure (Yeaman 2022) to explain when one or the other architecture will more likely emerge. However, most of these studies focus on scenarios where adaptation is assumed to occur, often explaining outcomes retrospectively in populations that, by definition, have adapted, which incurs a survival bias by not analyzing conditions when populations go extinct.

The question of how within-species variation and different genetic architectures may condition populations to rapidly adapt (evolutionary rescue) or become extinct has been more limited and has shown mixed conclusions. Gomulkiewicz and Holt theoretically demonstrated that evolutionary rescue should be possible under both single-locus and quantitative models, assuming high initial genetic variance and moderate selection. (Gomulkiewicz and Holt 1995). Building on this, Orr and Unckless (2008) argued that polygenic architectures should offer greater potential for rescue due to their genetic redundancy, allowing adaptive allele frequency shifts at many loci without relying on rare, large-effect mutations. However, subsequent work by Gomulkiewicz et al. demonstrated through analytical models and simulations that increasing the number of loci underlying fitness can dilute the strength of selection acting on each locus, thereby slowing the rate of adaptation and narrowing the conditions under which populations can persist following an environmental shift (Gomulkiewicz et al. 2010). More recently, Kardos and Luikart showed through deterministic models and stochastic simulations that polygenic trait architectures—by maintaining higher short-term evolutionary potential—can substantially improve population viability during rapid environmental change compared to traits controlled by few large-effect loci (Kardos and Luikart 2021).

To clarify this unresolved question, we conducted a large-scale set of forward-in-time population genetic simulations to test how genetic architecture influences the probability of evolutionary rescue under rapid environmental change within 10 generations. We modeled a quantitative trait under varying levels of polygenicity and heritability, and tracked population outcomes across different magnitudes of environmental change. By explicitly comparing polygenic and oligogenic architectures, we examined how genetic architecture shapes both the likelihood and predictability of evolutionary rescue. To ensure biological relevance, our simulations were grounded in empirical genomic data from *Arabidopsis thaliana*, a model system in eco-evolutionary dynamics (Leventhal, Ruffley, and Exposito-Alonso 2025).

## Materials and Methods

We focus on modeling rapid adaptation from standing genetic variation, as the pace of global change may require plant and animal species to rely on pre-existing diversity (Barrett and Schluter 2008). To simulate realistic genomic architectures, we used genome-wide variation data from a diverse founder population, which includes 10 individuals from 231 different *Arabidopsis thaliana* accessions variable in 3,235,480 SNPs, mimicking empirical experimental evolution in outdoor gardens of this species (GrENE-net.org project) (Wu et al. 2025). To load genome-wide variation in population genetic simulations with SLiM (Kelleher et al. 2018; Haller et al. 2019), we implemented a tree sequences approach (Kelleher et al. 2019), where neutral mutations were initially removed and represented by the ancestral recombination graph so only trait-relevant mutations were monitored through the simulations.

We modeled a single adaptive trait under natural selection. The trait’s genetic architecture varied in three parameters: polygenicity, initial allele frequency, and effect size distribution. Polygenicity ranged across nine levels (1–1000 loci; **Fig. 1**). Initial allele frequencies were sampled from the *A. thaliana* site frequency spectrum. Effect sizes were either drawn from a normal distribution N≈(0, 2) or inversely proportional to allele frequency, reflecting a trade-off between allele frequency and effect size (Simons et al. 2022). Since both approaches produced the same qualitative patterns of evolutionary rescue, we focus on the inverse model in the main results. We assume that allelic effects on the trait combine in an additive way: *A* _*n*_*=*∑*X*_*i*_*a*_*i*_, where *A* _*n*_ is the trait’s genetic value of the individual *n, X* is the individual’s genotype at the *i* _*th*_ contributing locus, and a _i_ is its effect size, and V _A_ is the variance in genetic values across our population. *A* _*n*_ were standardized in the first generation to ensure comparability across simulations, such that *μ = 0, σ*^*2*^ *= 1*. We simulated traits across five different heritability levels, h^2^ = V _A_ / V _A_+V_E_, ranging from 0.1 to 0.9. (**Fig. 1**), by creating environmental variance V _E_ such: *V* _*E*_ *= (V* _*A*_ *-h*^*2*^ *· V* _*A*_*) / h*^*2*^. Environmental noise for each individual was sampled as *en* _*n*_ taken from *N(0, V* _*E*_), and phenotypes were calculated as *Z* _*n*_ *= A* _*n*_ *+ en* _*n*_.

**Fig. 1.**
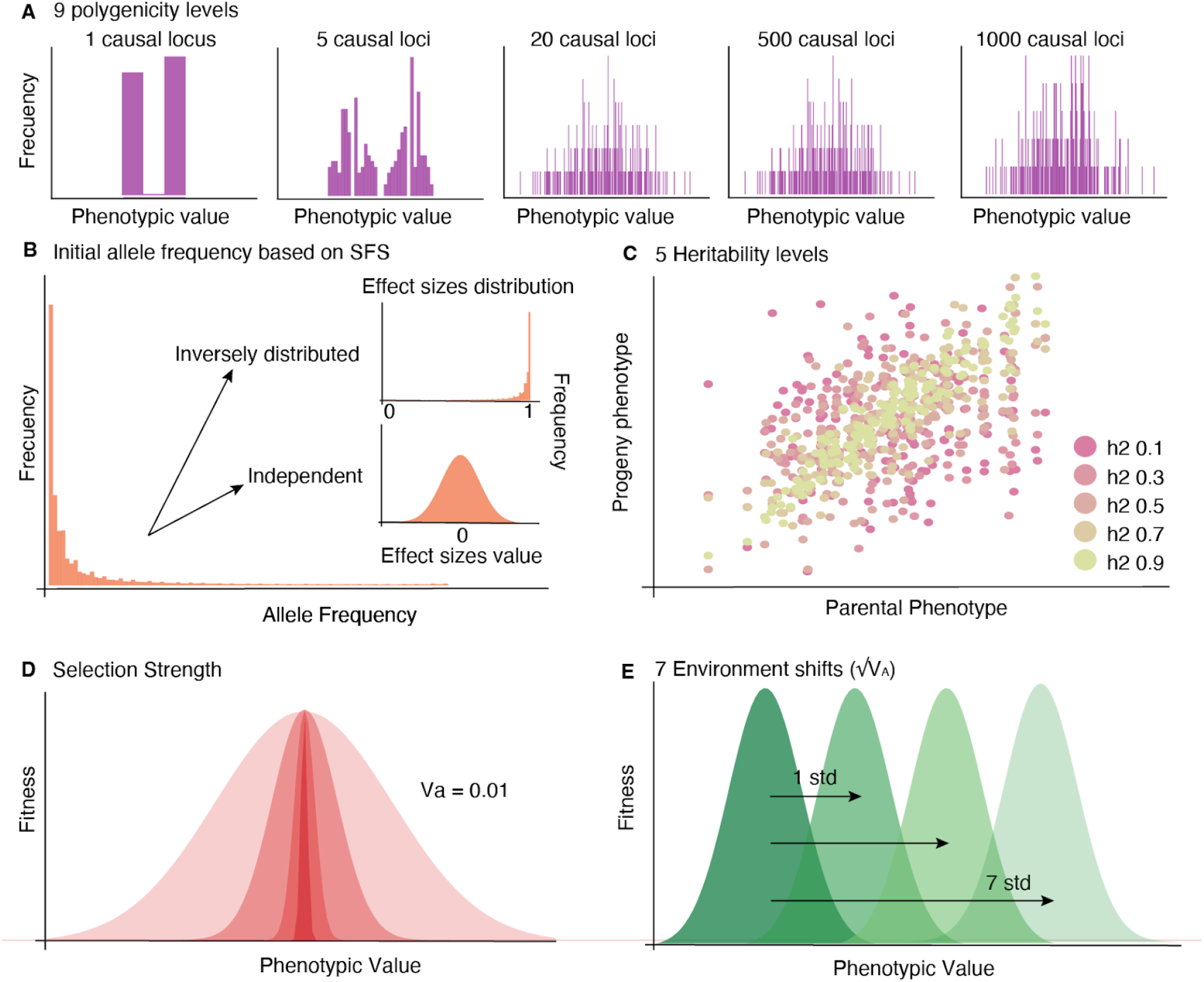
Parameter space. We varied **A**. the number of contributing loci [1,2,5,10,20,50,100,500,1000], **B**. initial allele frequency taken from the founder population site frequency spectrum, effect sizes of the contributing loci taken from a distribution inverse to the initial allele frequency (dependent) or from a normal distribution with mean 0 and standard deviation 2 (independent), **C**. heritability levels [0.1, 0.3, 0.5, 0.7, 0.9], **D**. selection strength modeled as fitness decay with Va=0.01, and **E**. distance to the new phenotypic optimum modeled as standard deviations from initial phenotypic mean [0, 1, 2, 3, 4, 5, 6, 7].

To model environmental change, we simulated eight levels of phenotypic optimum shifts (i.e. environmental change), measured in standard deviations from the initial mean *A* _*n*_ (**Fig. 1**). Viability selection was applied via the stabilizing selection model where *Fitness* _*n*_ *= exp(− (0*.*5 × (Z* _*n*_ − *Z* _*opt*_)^*2*^*) / V* _*s*_ ) and *Z* _*opt*_ is the phenotypic optimum and *V* _*s*_ represents selection strength. We conducted simulations at different magnitudes of environmental shift: Δ _*shift*_*=Z* _*avg*_ *-Z* _*opt*_; which we measure in units of starting population phenotypic standard deviation √*V* _*A*_. We initially simulated varying selection strengths, but were set to *V* _*s*_ = 0.01. We focused on scenarios with both strong selection (*V* _*s*_), and large-magnitude environmental shifts (√*V* _*A*_) to generate conditions where populations would either adapt or go extinct. This combination was necessary to produce classic evolutionary rescue dynamics, in which a sudden and substantial environmental shift causes an initial decline in population size, followed by either recovery or collapse (Fig. 5C) (note that this corresponds to dynamics of environmental shifts beyond what Milligan et al. studied (2025).

To conduct simulations that capture *A. thaliana* biology, we conducted simulations with a population size of 2310 individuals and were limited to a carrying capacity of 900 at the end of each generation, based on GrENE-net field garden observations (Wu et al. 2025). Generations were annual, non-overlapping, and assumed a 97% selfing rate (Platt et al. 2010). Offspring production followed a Poisson distribution with a mean of 7.247, based on experimental field data (Exposito-Alonso lab, unpublished). We incorporated a mean recombination rate of ≈4 cM/Mb (Singer et al. 2006).

For each level of polygenicity (×9), we generated 30 independent genetic architectures (×30), which were combined with five heritability levels (×5) and seven environmental shift scenarios (×7). Each unique combination was replicated five times (×5) to account for stochasticity in the simulations. In total, we performed 47,250 forward-in-time genetic simulations, each of them tracking a single population for 10 generations. At each generation, we extracted key data including population size, pseudo-environmental variables, individual genotypes, phenotypes, and fitness values, in addition to saving full tree sequences for downstream analysis.

Simulations were implemented in a reproducible Snakemake pipeline (Mölder et al. 2021)https://github.com/Tatianabellagio/slim_grenenet/tree/es_dep_af.(Mölder et al. 2021), which takes the starting VCF, runs all simulation combinations, and produces summary outputs.

## Results

### Simulation parameters defining population mortality versus evolutionary rescue

To examine how heritability, polygenicity, and the magnitude of environmental change affect the ability of a population to adapt and persist, we first performed a multiple logistic regression across all simulations. As expected, survival probability declined sharply with increasing environmental change (i.e., distance shift of the phenotypic optimum, *z* = -92, *P* < 1×10^−2^, *n* = 43200), while heritability had a strong positive effect on survival (*t* = 80, *P* < 1×10^−2^, *n* = 43200). Notably, polygenicity also had a small but statistically significant positive effect (*z* = 23, *P* < 1 × 10^−2^, *n* = 43200).

Given the modest effect size of polygenicity in population survival, we hypothesized that the influence of polygenicity on survival might be context-dependent. To explore this, we ran independent logistic regressions within fixed combinations of polygenicity, heritability, and environmental shift, allowing us to isolate the effect of a single parameter on survival while holding all others constant—such that variation across replicates could be attributed to that parameter and to stochasticity from selection and drift. This generated various effect sizes for each predictor across contexts (**Fig. 2**, only predictors with Bonferroni-corrected *P-value* < 0.05 were retained). This approach confirmed the consistent positive role of heritability (*z* = 3.9–13.3, *P* <9×10^−05^, n=40), and the strongly negative impact of environmental change (*z* = -18.7 – -9.6, *P* < 8×10^−22^, *n*=27). However, the relationship of polygenicity with population rescue spanned both positive and negative ranges (*z* = -4.8 – -17.4, *P* < 9×10^−4^, *n*=17), confirming that the effect of polygenicity on survival depends on the specific selection and genetic context.

**Fig. 2.**
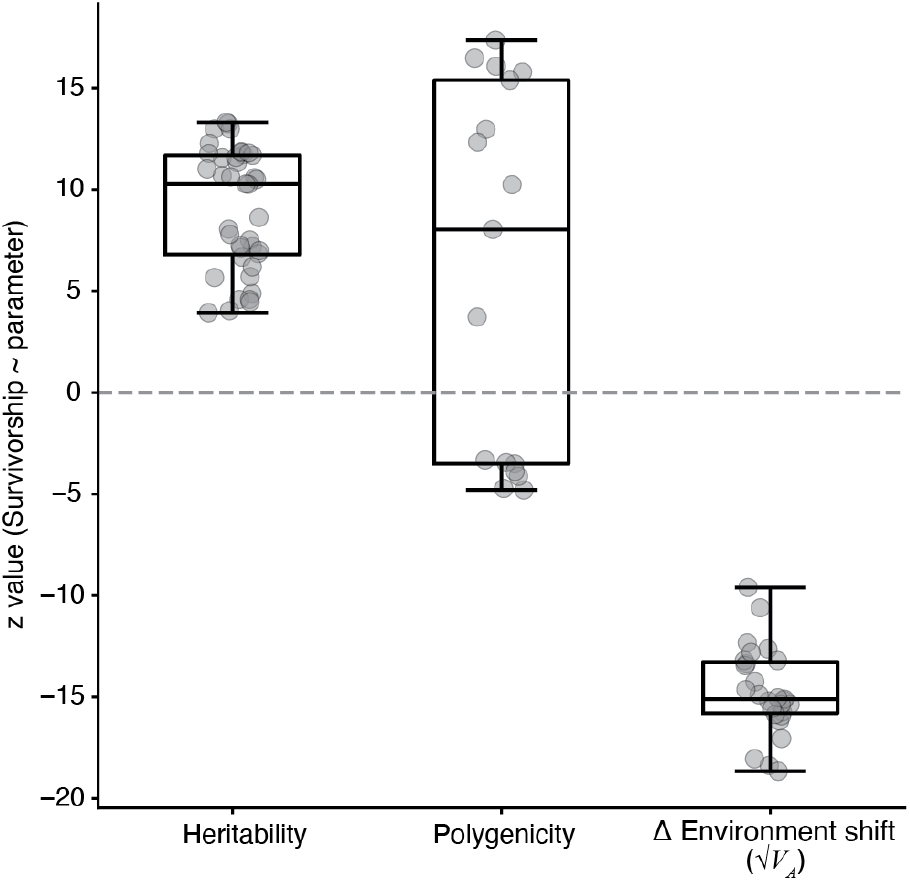
Distribution of t-values for logistic regressions on survival. Logistic regressions were performed across fixed combinations of other simulation parameters. Each bar shows the parameter’s impact when all other variables were held constant. Only significant (Bonferroni-adjusted) results are shown.

### Polygenicity has context-dependent effects on evolutionary rescue

We further explored the combination of parameters that contributed to either a positive or negative effect of polygenicity on population survival, and observed that increased polygenicity had a positive impact on population survival under low-magnitude environmental changes (Δ _*shift*_ √*V* _*A*_ to new optimum=0–3; *z* = 3.7–17.4, *P* < 2×10^−04^) where the overall population survivorship was high (38–99%). Conversely, the opposite effect was observed under large environmental changes (Δ _*shift*_=5–7; *z*=-4.8 – -3.3, *P*< 9 ×10^−04^) where low-polygenic architectures led to increased survival, although overall population survival was extremely low (1-9%) (**Fig. 3**).

**Fig. 3.**
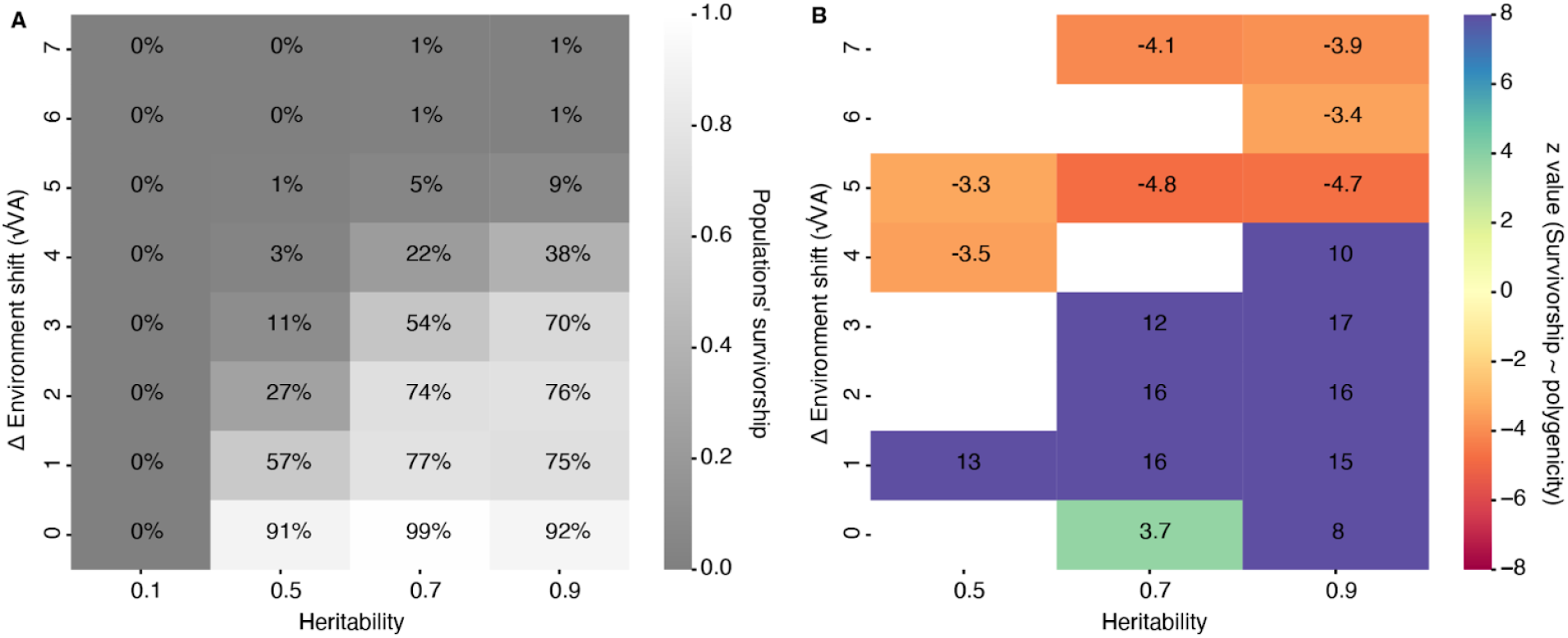
Impact of polygenicity on population survival after the 10^th^ generation. A. Survival rates across combinations of heritability and environmental shift. **B**. *z values* from logistic regressions showing the effect of polygenicity on survival, for each parameter combination.

### Fitness variance heterogeneity as a predictor of population survival

We observed that genetic variance in fitness, *V* _*w*_, varied substantially across replicates, even when simulations shared the same genetic architecture. Although all architectures started with fixed mean genetic value *A* and genetic variance *V* _*A*_, randomness in the sampled allelic effects and number of contributing loci introduced differences in *V* _*w*_. Supporting the long-standing theory that *V*_*w*_ reflects adaptive potential (Fisher 1930), we found a strong positive relationship between initial *V*_*w*_ and population survival across all scenarios (*z*=18.47, *P*<1×10^−2^, holding h^2^=0.7 constant, **Fig 4**). To better understand the context-dependent effects of polygenicity on survival identified earlier, we then explored how polygenicity influences the mean and variability of *V* _*w*_ under a gradient of Δ _*shift*_ environmental change (**Fig. 5A**, constant h^2^=0.7). Across this gradient, we observed a consistent pattern: under mild environmental change *V* _*w*_ increased significantly with polygenicity (Δ_*shift*_ < 2, *P*<1×10^−2^), supporting higher survival probabilities in highly polygenic architectures. Under large environmental changes, the relationship inverted: higher average *V*_*w*_ was associated with lower polygenicity (Δ_*shift*_ > 3, *P*<1×10^−2^). This pattern helps explain why low-polygenic architectures sometimes outperformed, on average, highly polygenic ones under extreme environmental shifts, despite greater variance in survival outcomes.

**Fig. 4.**
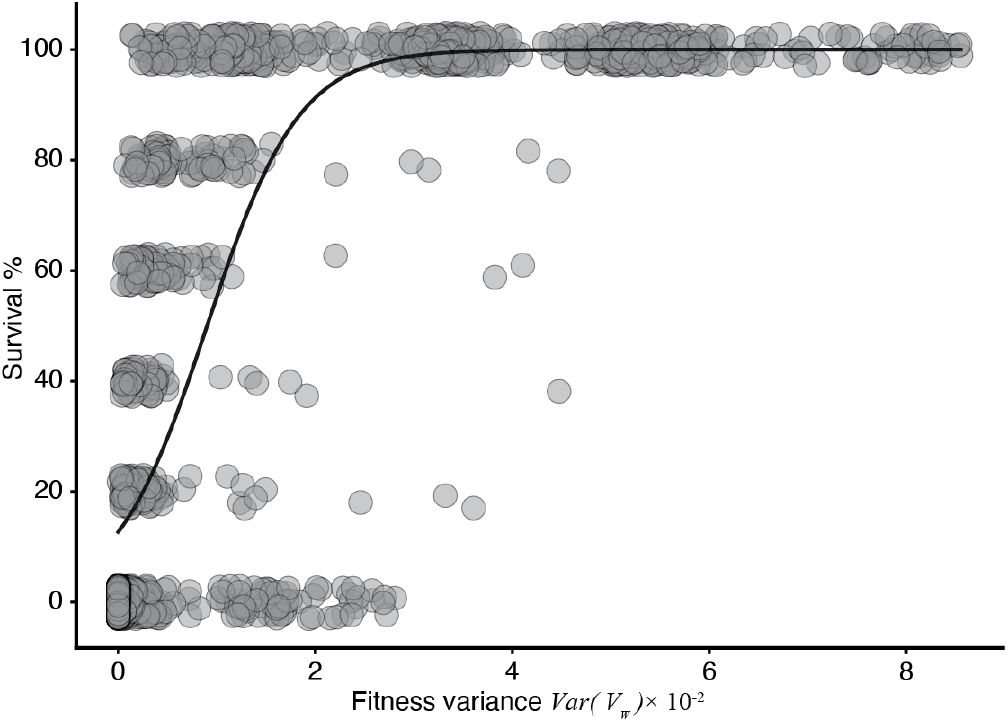
Relationship between initial additive genetic variance in fitness *V*_*w*_ and population survival on the last generation. Each point represents a unique parameter combination with survival expressed as the percentage of replicates (out of 5) in which the population persisted. Heritability was held constant at h^2^ = 0.7.

**Fig. 5.**
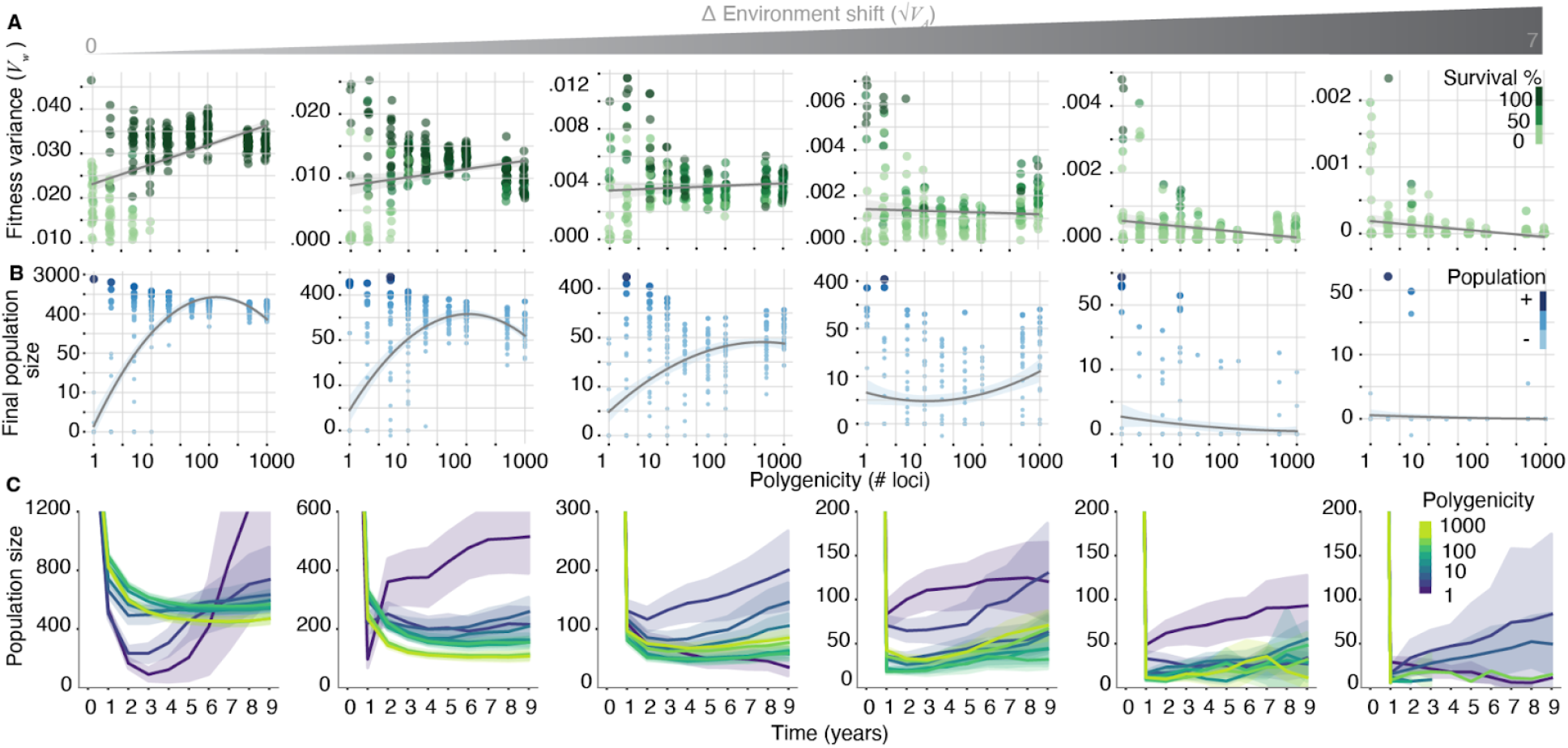
Effects of polygenicity on population genetics and demography under varying magnitudes of environmental change. Heritability was held constant at h^2^ = 0.7. **A**. The effect of polygenicity on additive genetic variance in fitness (*V*_*w*_) depends on the magnitudes of environmental change. Under small environmental shifts (1-2 SD), *V* _*w*_ increased significantly with greater polygenicity (*P* < 0.001). This relationship weakened and became nonsignificant under intermediate shifts (3-4 SD, *P* = 0.1881), and reversed under large shifts (5-6 SD), where lower-polygenic architectures showed higher *V* _*w*_ (*P* < 0.001). **B**. Relationship between polygenicity and final population size (10^th^ generation) under different magnitudes of environmental change. Low-polygenic architectures exhibited greater variance in final population size across all magnitudes of environmental change, with some replicates achieving the largest population sizes. High-polygenic architectures showed more consistent demographic outcomes that closely mirrored survival probabilities. **C**. Population trajectories across time under different polygenic architectures and environmental shift magnitudes. All simulations began with an initial population size of 2,310. When survival occurred, populations typically exhibited U-shaped demographic trajectories characteristic of evolutionary rescue.

To further explore how polygenicity influences demographic outcomes, we examined both the final population sizes (**Fig. 5B**) and temporal trajectories (**Fig. 5C**) across simulation replicates. Despite the overall trend that low-polygenic architectures were associated with lower survival probabilities, we observed that, conditional on survival, monogenic architectures led to larger final population sizes compared to high-polygenic populations across all levels of environmental change; this is likely due to the dynamics of adaptation being more rapid. Furthermore, low-polygenic populations exhibited greater variability in final size, likely driven by high variable fitness variance (*V* _*w*_). In contrast, high-polygenic architectures showed much less variability: population sizes tightly mirrored survival trends, with stable populations under small environmental shifts and sharp declines toward extinction under severe shifts.

To explain the counterintuitive relationship of *V* _*w*_ and polygenicity, we studied the variation in *V*_*w*_ across simulations. We found that the variability of *V*_*w*_ across replicates was consistently greater in low-polygenic architectures, regardless of the magnitude of environmental change. Using multiple regression while controlling for heritability and environmental shift, we confirmed that lower polygenicity was associated with significantly higher variability *sd(V* _*w*_) across simulation replicates (polygenicity *t* =-5.496, *P* <1×10^−2^) and higher skew under moderate environmental change (*Skew[V* _*w*_] _*oligo*_/*Skew[V* _*w*_ ] _*poly*_= 2.1). Furthermore, the variability in *V* _*w*_ across simulations decayed rapidly around 30 contributing loci (**Fig. 6**), suggesting a potential threshold separating architectures with stochastic versus more predictable evolutionary dynamics, coinciding with the central limit.

**Fig. 6.**
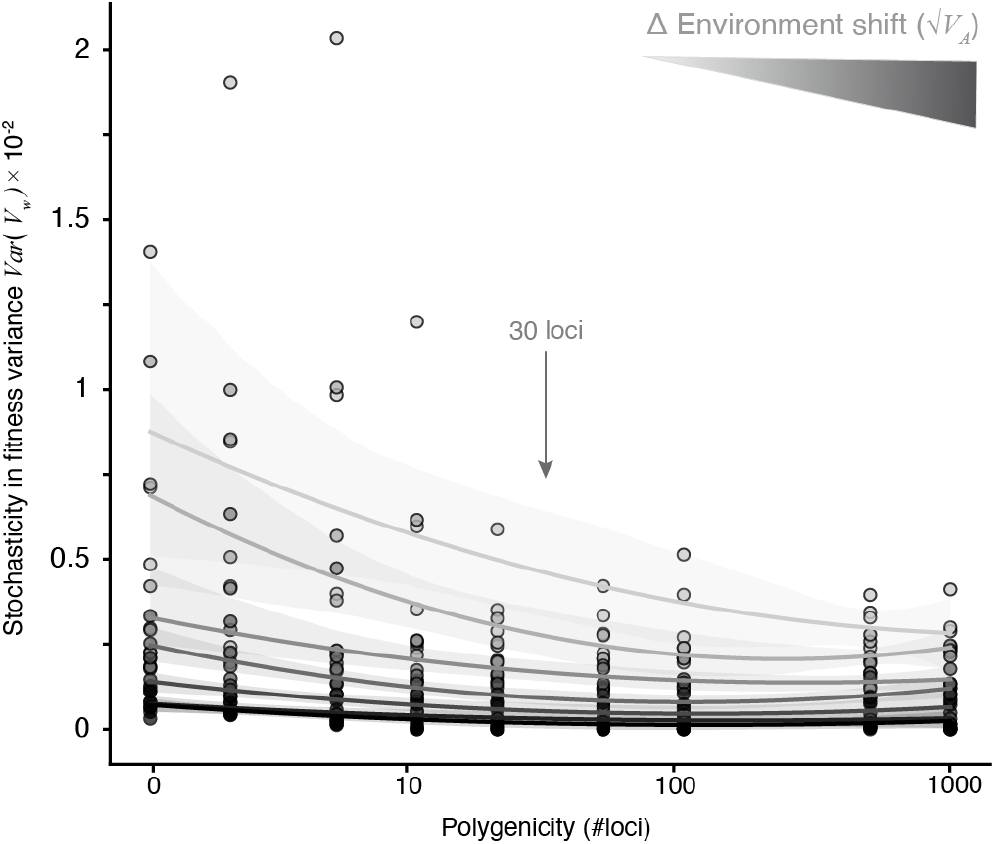
Relationship between polygenicity and stochasticity in fitness variance (V _w_) across simulations under different magnitudes of environmental change. Lower-polygenic architectures exhibit significantly greater variability in V _w_ compared to highly polygenic ones. A sharp decline in V _w_ variability occurs around log(30) contributing loci.

### Predictability of evolutionary rescue under monogenic and polygenic architectures

Building on the observation that fitness variance, *V* _*w*_, is more volatile under low-polygenic architectures than polygenic architectures (**Fig. 6**), we asked whether this variability could impact the predictability of evolutionary rescue. To investigate this, we categorized simulations into low- and high-polygenic groups based on the number of contributing loci, using 30 loci as a threshold identified from the decay in V _w_ variability. Within each group, we trained a decision tree classifier (depth = 2) using a 70-30 train-test split, with heritability, polygenicity, and environmental shift magnitude as predictors (**Fig. 7**). For high-polygenic architectures (>30 loci), survival was highly predictable: the decision tree correctly classified 91% of simulations in the test set. In this model, the primary predictor of survival was the magnitude of environmental shift (feature importance = 0.52), with a threshold at 3.5 √*V* _*A*_ separating surviving and non-surviving populations. In contrast, for low-polygenic architectures (<30 loci), survival was unpredictable. In fact, the model always predicted extinction under monogenic architectures (i.e., this was often successful since monogenic architecture simulations led to 84% extinction; the model, however, recalled zero survival cases. This indicates that the model did not find useful heritability and or environmental shift information as predictors. In contrast, in high-polygenic architectures, heritability was the second most important predictor (threshold *h*^*2*^ = 0.3), while polygenicity itself (very large vs moderately large number of loci) had no predictive value. Overall, these findings suggest that low-polygenic architectures generate non-predictable survival outcomes. These findings support the existence of a tipping point near 30 contributing loci, beyond which evolutionary outcomes become more deterministic and less sensitive to the underlying genetic architecture, including stochastic differences in initial allele frequencies and effect sizes.

**Fig. 7.**
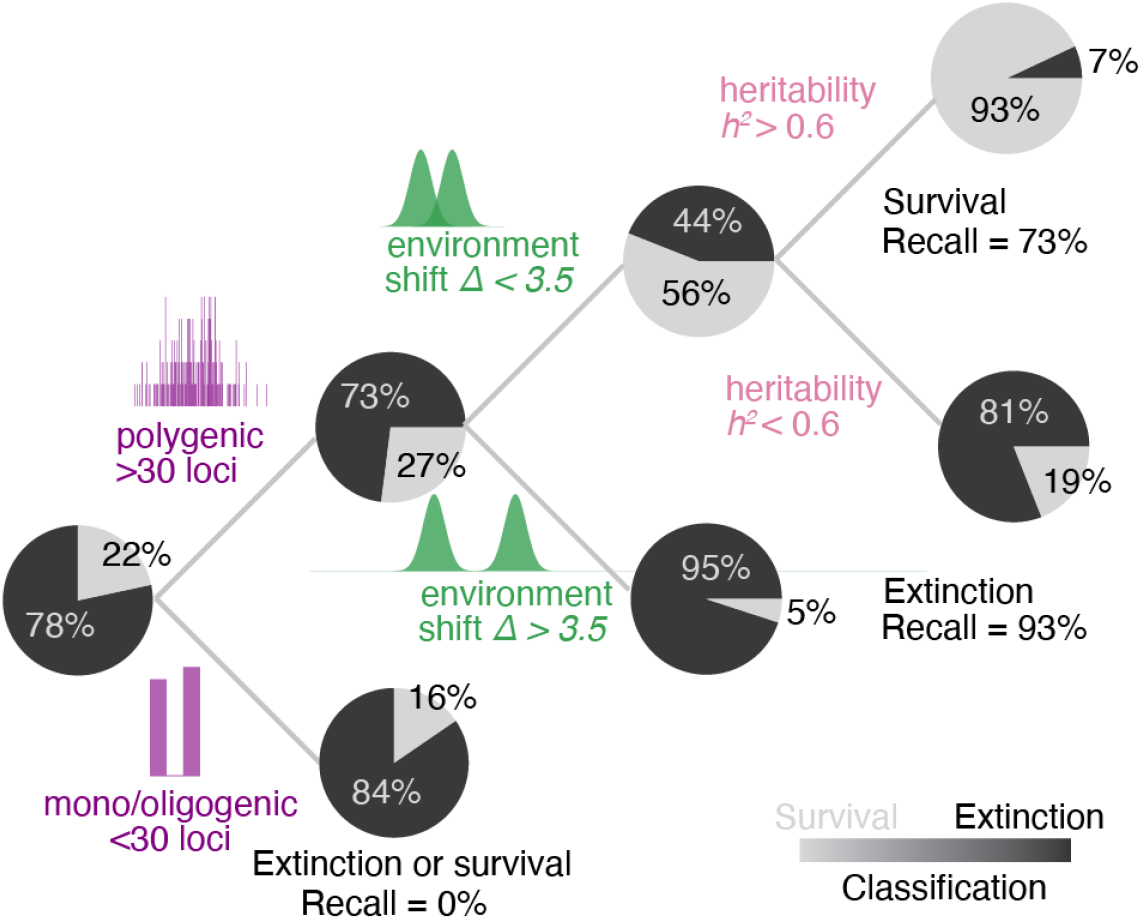
Decision tree models trained to predict population survival based on heritability, polygenicity, and the magnitude of environmental change. Two separate models were trained: one for low-polygenic architectures (<30 loci) and one for high-polygenic architectures (>30 loci). At each node, pie charts show the proportion of simulations classified as survived or extinct. Branch splits reflect thresholds learned by the model during training.

## Discussion

In nature, there are abundant examples of adaptation occurring either through polygenic or monogenic architectures (Bomblies and Peichel 2022), a distinction that may often be idiosyncratic to the ecology of species and its environment. Perhaps then the key question in the longstanding “many vs few genes” debate is not whether polygenic or monogenic architectures may be more or less common, but rather under which environmental conditions each genetic architecture is most likely to fuel rapid evolutionary rescue. Our key result is that the effectiveness of a given genetic architecture in promoting evolutionary rescue depends critically on the magnitude of environmental change, with polygenic architectures favored under milder shifts, and monogenic architectures under more severe ones.

Recent studies have begun to clarify the ecological and genetic factors that influence whether adaptation is driven by a few genes of large effect or many genes of small effect. Höllinger et al. emphasized that monogenic and polygenic architectures are all but the same with the key distinction being simply the amount of background loci in the genome that redundantly provide similar paths to adaptation (Höllinger, Pennings, and Hermisson 2019). Yeaman et al. further showed that the type of selection acting on a trait—directional vs spatially-varying—can influence the emergence of different architectures, with spatially-varying selection tending to favor fewer, larger-effect loci, and directional selection favoring more polygenic responses (Yeaman 2022). Most recently, Milligan et al. showed that polygenic adaptation tends to dominate under moderate environmental shifts, while large-effect alleles are more likely to fix when shifts are larger. However, this study is limited to environmental changes no greater than the width of the fitness function (√*V*_*s*_), explicitly excluding scenarios where the shift is severe enough to risk extinction. Within this bounded regime, they focus on allele dynamics assuming constant population size, and show that large-effect alleles are often selected against during the stabilizing selection phase after the early directional selection phase.

In contrast, fewer studies have explicitly examined the effect of genetic architecture in scenarios where populations would go extinct in the absence of rapid adaptation, that is, within the framework of evolutionary rescue. Gomulkiewicz et al. modeled adaptation to harsh environments and showed that a combination of a rare major-effect allele and polygenic background variation can enhance adaptation in a new environment (Gomulkiewicz et al. 2010). Their model predicted that adaptation driven solely by diffuse, small-effect loci may be too slow to enable persistence under strong selective shifts. Our results partially support this view: under severe environmental change, high-polygenic architectures alone often failed to produce the extreme phenotypes required for rescue (i.e. the phenotypic distribution did not overlap with the new fitness distribution), reinforcing the idea that polygenic architectures may have limited evolutionary potential. However, we think their model outcome is partly driven by their assumption of fixed *V* _*w*_ across genetic architectures at simulation onset, which can artificially dilute selection on polygenic traits. In contrast, our simulations allow trait distributions—and thus V _w_—to emerge organically from the underlying genetic architecture. Another later key study showed that polygenic architectures confer greater population viability during rapid environmental change, primarily due to their higher short-term evolutionary potential, which they attribute to a more stable maintenance of heritability (Kardos and Luikart 2021). Our findings align with this under mild to moderate environmental shifts, where polygenic traits consistently enhanced survival. Their definition of a new environmental optimum, 3.2 √*V* _*A*_ from the initial phenotypic mean, falls within what we classify as small to intermediate change. In those scenarios we also observe better polygenic performance. Notably, Kardos and Luikart also observed high variability across low-polygenic architectures and emphasized that their outcomes were sensitive to initial allele frequencies, an observation that closely mirrors our own.

A key insight of our findings is that the environment interacts with the genetic architecture to create unexpected distributions of genetic variances in fitness. Following Fisher’s Fundamental Theorem of Natural Selection (FTNS), which equates *V* _*w*_ to the rate of adaptation, we found that average *V* _*w*_ positively correlated with the probability of evolutionary rescue across all our simulations (**Fig 4**) (Fisher 1930). A fundamental observation from Fisher that led to integrating Mendelian genetics and biometrician views of inheritance was that as the number of mendelian loci involved in a trait increases, the distribution of phenotypes becomes a Gaussian distribution with population variance *V* _*A*_ (Fisher 1918). An obvious observation from first principles, and with critical importance in our study, is that genetic architectures of low numbers of loci (i.e., < 30) generate phenotypic distributions that do not conform to Gaussian assumptions. In our study we realized that non-normality of phenotypic distribution has critical consequences for fitness variance. Even when simulations started with the same phenotypic mean and variance (*V* _*A*_), the resulting distribution of *w* differed systematically across architectures (**Fig. 5, Fig. 6**). Low-polygenic architectures produced *V* _*w*_ with higher variance across replicated simulations (*Var[V* _*w*_] _*monogenic*_ *> Var[V* _*w*_ ] _*polygenic*_) and with significant skewness (*Skew[V* _*w*_ ] _*monogenic*_ *> 0, Skew[V* _*w*_ ] _*polygenic*_ *∼ 0*), leading to more volatile and unpredictable evolutionary rescue outcomes. Furthermore, the average *V* _*w*_ varied across environments such that under small environmental change polygenic architectures led to on average higher (*E[V* _*w*_ |Δ _*small*_ ] _*polygenic*_ *> E[V* _*w*_ | Δ _*small*_ ] _*monogenic*_) while the opposite happen under more extreme environments *E[V* _*w*_|Δ _*big*_ ] _*monogenic*_ *> E[V* _*w*_ |Δ _*big*_ ] _*polygenic*_). The fact that previous studies fixed *V* _*w*_across architectures, prevented studying the interaction of environment and architecture shaping genetic variance. We hence suggest studying this phenomenon analytically in future work.

Contemporary conservation genetics has mainly focused on protecting genetic diversity within animal and plant species, to indirectly protect their long-term evolutionary potential (Allendorf, Luikart, and Aitken 2012). More recently, there have been calls to include genomics as it may help anticipate (mal)adaptive conditions of populations based on their mismatch of locally adaptive genetic variation with future climate shifts (Capblancq et al. 2020). Our results suggest a key dimension to understanding the (un)predictability of future evolutionary rescue is the genetic architecture. Some recent studies have attempted to quantify, for instance, the heritability *H* _*w*_^*2*^ and *V* _*w*_ of fitness through pedigrees in wild populations to quantitatively assess adaptation potential. For instance, Bonnet et al. (2022) found *V* _*w*_ values significantly different from zero in 10 of 19 vertebrate natural populations, although the ratio to total fitness variance was small, *h*^*2*^_*w*_ *=* 3% (Bonnet et al. 2022). In plant common gardens in herbs and provenance trials of trees, where genotypes from different source populations are compared together in the same environment, often show significantly larger heritabilities in fitness traits: *h*^*2*^ *∼* 0-0.64 (Roybal and Butterfield 2018; Bemmels and Anderson 2019; Leventhal, Ruffley, and Exposito-Alonso 2025). In *A. thaliana* common gardens, where fitness is often directly measured and used for GWAs, previous studies have found a highly polygenic architecture (Exposito-Alonso et al. 2019; Leventhal, Ruffley, and Exposito-Alonso 2025; Wu et al. 2025). Common garden studies, however, do not reflect the capacity of local natural populations to adapt. A recent experimental evolution study conducted in outdoor plots to investigate rapid adaptation and extinction in nature inspired our simulations (Wu et al. 2025). A key result of this experimental evolution study is the interaction of *V* _*w*_, the transplant environment, and long-term population survival (**Fig. 8**). Under extreme environmental shifts—such as the transplant distance of *A. thaliana* populations to a common garden measured based on temperature of origin—a high *V* _*w*_ measured in the first year of the experiment was predictive of survival within 5 years. Such experimental approaches modifying the starting genetic composition of the population would be ideal to better understand the complexities of genetic architectures and adaptation in realistic environments.

**Fig. 8.**
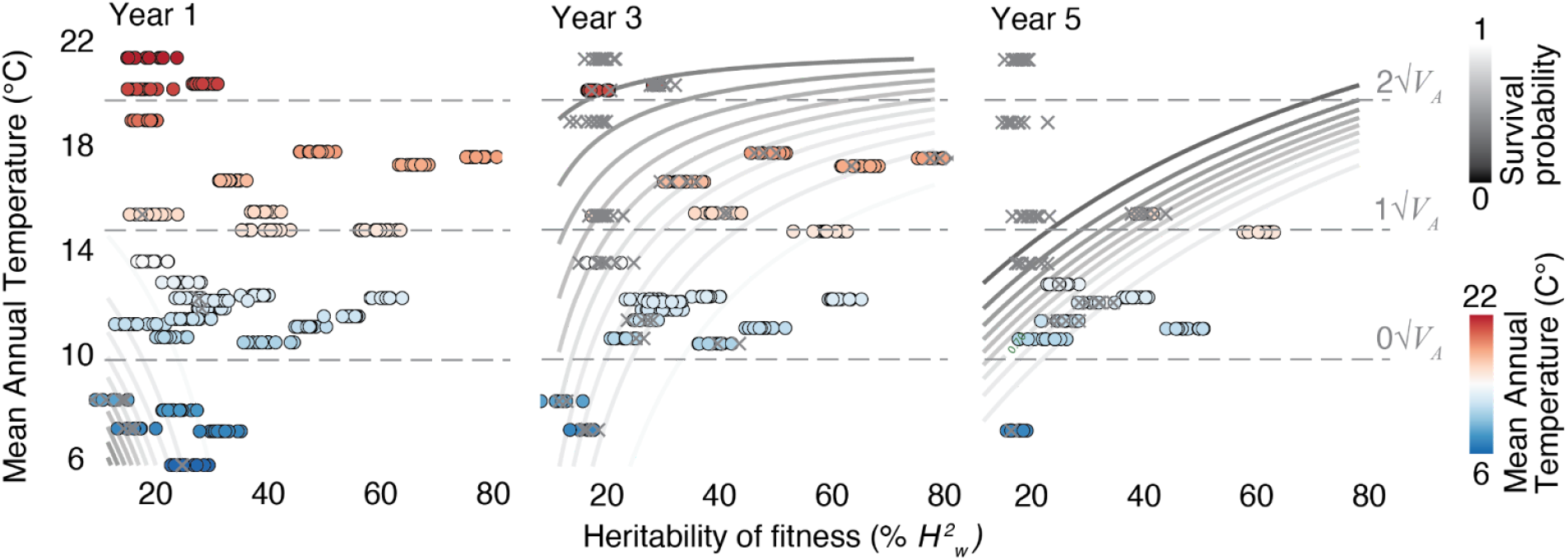
Interaction between environment and genetic variance in fitness in experimental evolution plots of *A. thaliana*. Modified from (Wu et al. 2025). *Arabidopsis thaliana* populations were planted in field sites spanning a climate gradient. Circles represent surviving populations, and Xs denote extinction. Survival probability was modeled using ecotype-composition predictability across environments and climate (mean annual temperature). Dashed horizontal lines denote the climate expressed as √*V* _*A*_ of the ecotype’s climatic origin. X-axis values are jittered for clarity; each site has a single heritability estimate.

Incorporating the understanding of genomic architecture’s influence in evolutionary rescue into conservation management could provide valuable insights. For example, the efficiency of assisted gene flow may be affected by the genetic architecture of adaptation in a species. Simulations have shown that adaptation can occur rapidly following the translocation of individuals carrying one or a few large-effect alleles (Grummer et al. 2022). In contrast, when adaptation depends on many small-effect alleles, the benefits may take much longer to manifest, as each allele contributes only a small portion of the adaptive response and thus takes longer to introgress into the local population (Grummer et al. 2022). Furthermore, as genomic technologies become more accessible in non-model species, it has become increasingly clear that local adaptation often involves clusters of tightly linked loci—blurring the line between monogenic and polygenic architectures (Schwander, Libbrecht, and Keller 2014; Gutiérrez-Valencia et al. 2021). These so-called “supergenes” underlie examples of color polymorphisms in butterflies (M. J. Thompson and Jiggins 2014) and ecotypic differentiation in sunflowers (Todesco et al. 2020). Although these traits are genetically complex, due to tight linkage, their evolutionary dynamics better approximate monogenic adaptation (Oomen, Kuparinen, and Hutchings 2020; Yeaman 2022).

In principle, with perfect knowledge of the genetic basis and architecture of adaptation, it may be even possible to directly engineer adaptive traits through gene editing (Wang and Doudna 2023). This may be a last resort when within-species genetic diversity has been severely depleted—a scenario that is already a concern for many species (Exposito-Alonso et al. 2022). However, conservationists have rightly cautioned that uncertainty in identifying the correct traits or targets for intervention may result in ineffective or even harmful outcomes (Kardos and Shafer 2018). While current CRISPR/Cas9 technologies have advanced to the point that nearly *any* gene can be edited in plants and animals (Gilbertson, Puchta, and Slotkin 2025; Wang and Doudna 2023), the challenge lies in identifying which genes to target. As a result, gene editing efforts have so far focused on a few well-characterized traits—such as coat color in mice, recently modified to mimic that of woolly mammoths (Chen et al. 2025), or spring flowering time pathways in Brassicaceae, used to accelerate phenology in wild accessions (Ruffley et al. 2024; Lutz et al. 2024). Returning to the model system *Arabidopsis thaliana*, while fitness traits appeared to be polygenic in previous studies (Exposito-Alonso et al. 2019; Leventhal, Ruffley, and Exposito-Alonso 2025; Wu et al. 2025) specific key adaptive traits such as spring flowering time may be controlled by a few modifiable loci. For instance, targeted knock-out or knock-down of the *FLOWERING LOCUS C* (*FLC)* gene led to significantly earlier flowering consistently in 94 wild-type accession lines collected from different natural populations (≈60 days, **Fig. 9**). We estimated that such phenotypic shift corresponds to over >2√*V*_*A*_. Under these circumstances, for instance, if adaptation to global change was solely determined by accelerating flowering time, a large-effect loss-of-function *flc* mutation may catalyze evolutionary rescue. Yet this strategy becomes far more precarious in natural population conservation, where fitness is rarely determined by a single trait but rather emerges from the combined effect of many underlying traits, or where large-effect loci may be purged due to environmentally varying selection and stabilizing selection. As a result, fitness-related traits are typically highly polygenic, and their expression is further modulated by gene-by-environment interactions. From these principles, one may derive that gene editing could be used for genetic diversity generation for polygenic architectures, although such polygenic, complex genomic and chromosome rearrangement are still a frontier of technological development (Gilbertson, Puchta, and Slotkin 2025, Exposito-Alonso *in preparation*).

**Fig. 9.**
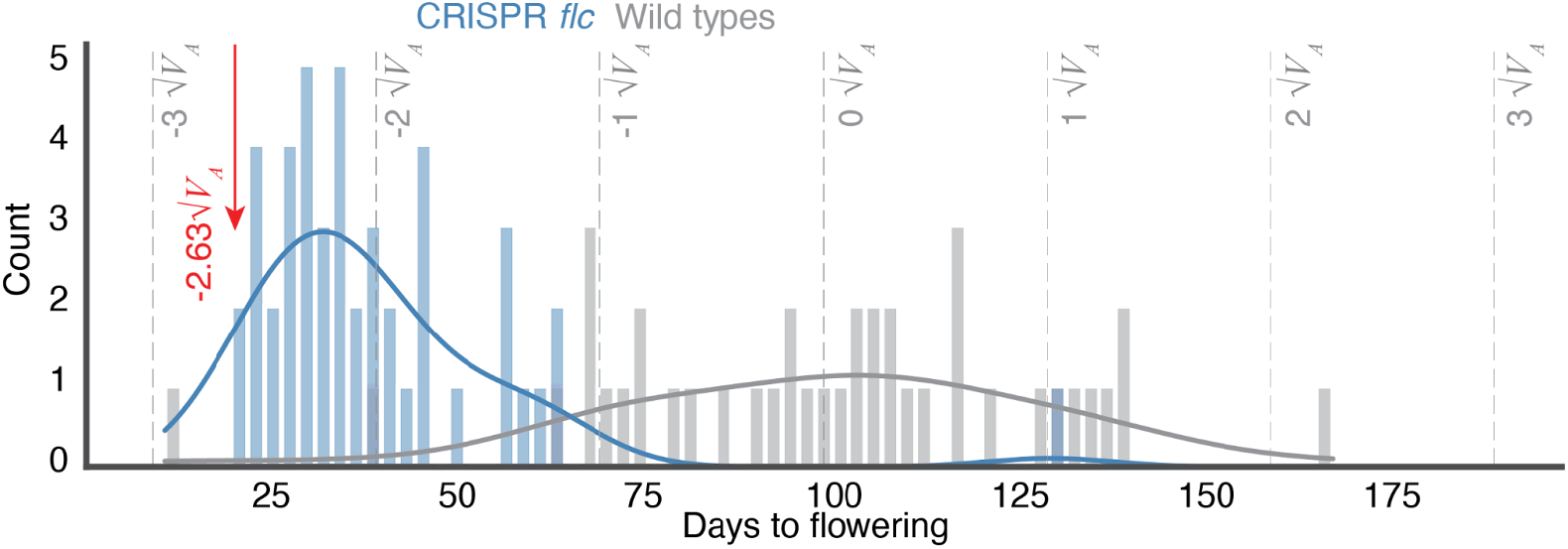
Distribution of mean flowering time per ecotype for wild-type (wt) and CRISPR *flc* knockout (KO) individuals across 94 *Arabidopsis* ecotypes. Modified from (Ruffley et al. 2024). For each genotype, flowering time was averaged across three replicates. Vertical dashed lines show √*V*_*A*_ of the wild-type distribution. The red arrow marks the minimum flowering time observed among the flc KO lines, which lies 2.63 √*V* _*A*_ units below the wt mean. All plants were grown under long-day (LD) conditions at 22 °C.

As anthropogenic environmental change accelerates, accurately predicting the probability, pace, and direction of evolutionary rescue is critical for biodiversity conservation and ecosystem management (Schoener 2011; Bay et al. 2017; Coulson et al. 2017; Forester et al. 2022). Advances in genomic technologies now make it possible to estimate both the adaptive potential and the underlying genetic architecture of adaptive traits in natural populations, providing critical information for improving predictive models of population persistence and guiding conservation strategies (Shafer et al. 2015; Hoffmann et al. 2015; Funk et al. 2012; Allendorf, Luikart, and Aitken 2012; McLachlan, Hellmann, and Schwartz 2007; Thomas et al. 2013). Our results highlight the value of incorporating genetic architecture parameters into predictive frameworks, as they significantly influence both the likelihood and predictability of adaptive responses. In particular, populations where adaptation will depend on highly polygenic traits may offer predictability and greater resilience to moderate environmental shifts, while those depending on low-polygenic adaptive traits may occasionally achieve rescue under extreme change, albeit with higher stochastic risk.

## Acknowledgements

We are thankful for feedback and discussions with Lucas Czech, Xing Wu, and other members of the MOILAB. We thank Ben Haller for feedback on SLiM code and Peter Ralph and Clemens Weiß for feedback on tree sequence implementation.

## Author contribution

T.B. and M.E.-A. designed the project. T.B. wrote the code, conducted analyses, and prepared the manuscript draft. T.B. and M.E.-A. wrote and edited the final manuscript.

## Funding statement

M.E.-A. is supported by the Office of the Director of the National Institutes of Health’s Early Investigator Award (1DP5OD029506-01), the U.S. Department of Energy, Office of Biological and Environmental Research (DE-SC0021286), by the U.S. National Science Foundation’s DBI Biology Integration Institute WALII (Water and Life Interface Institute, 2213983), by the Carnegie Institution for Science, the Howard Hughes Medical Institute, and the University of California Berkeley.

Computational analyses were done on the High-Performance Computing clusters of the Carnegie Institution for Science and High Performance Computing cluster of the University of California Berkeley.

## Data and code availability

All code required to reproduce the analyses presented in this study is openly available at: https://github.com/Tatianabellagio/slim_grenenet/tree/es_dep_af

